# Depleting Hypothalamic Somatostatinergic Neurons Recapitulates Diabetic Phenotypes in Brain, Bone Marrow, Adipose, and Retina

**DOI:** 10.1101/2021.03.30.437706

**Authors:** Chao Huang, Robert Follett Rosencrans, Raluca Bugescu, Cristiano P. Vieira, Ping Hu, Yvonne Adu-Agyeiwaah, Karen L Gamble, Ana Leda F. Longhini, Patrick M Fuller, Gina M. Leinninger, Maria B. Grant

**Affiliations:** Department of Ophthalmology, University of Alabama at Birmingham, Birmingham, Al 35205; Department of Physiology, Michigan State University, East Lansing, MI 48824; Department of Psychiatry and Neurobehavioral Neurobiology, University of Alabama at Birmingham, Birmingham, Al 35205; Department of Neurology, Beth Israel Deaconess Medical Center and Division of Sleep Medicine, Harvard Medical School, Boston, MA 02115

**Keywords:** hypothalamus, somatostatin, diabetes, monocytosis, neuroimmunology, retina, electroretinogram

## Abstract

Hypothalamic inflammation and sympathetic nervous system hyperactivity are hallmark features of metabolic syndrome and type 2 diabetes. Hypothalamic inflammation may aggravate metabolic and immunologic pathologies due to extensive sympathetic activation of peripheral tissues. Loss of somatostatinergic (SST) neurons may contribute to enhanced hypothalamic inflammation. The present data show that leptin receptor deficient (db/db) mice exhibit reduced hypothalamic somatostatinergic cells, particularly in the periventricular nucleus. We model this finding, using adeno-associated virus (AAV) delivery of diphtheria toxin (DTA) driven by an SST-cre system to deplete these cells in SST^cre/gfp^ mice (SST-DTA). SST-DTA mice exhibit enhanced hypothalamic c-fos expression and brain inflammation as demonstrated by microglial and astrocytic activation. Bone marrow from SST-DTA mice undergoes skewed hematopoiesis, generating excess granulocyte-monocyte precursors and increased pro-inflammatory (CCR2^hi^) monocytes. Visceral adipose tissue from DTA-treated animals was resistant to catecholamine induced lipolysis. Finally, SST-DTA mice exhibited a “diabetic retinopathy like” phenotype: reduced visual function by optokinetic response and electroretinogram, as well as increased percentages of retinal monocytes. Importantly, hyperglycemia was not observed in SST-DTA mice. Thus, the isolated reduction in hypothalamic somatostatinergic neurons was able to recapitulate several hallmark features of type 2 diabetes in disease relevant tissues.

---

Hypothalamic inflammation and sympathetic nerve hyperactivity are important upstream causes of metabolic and immunologic pathologies in diabetes and metabolic syndrome (METS). Both genetic and environmentally induced models of METS elicit hypothalamic inflammation and elevations in sympathetic outflow, often very early in the disease course[1–3]. Emerging evidence shows that local inflammation also sensitizes the hypothalamus to ascending sensory innervation, enhancing sympathetic outflow[4]. Thus, inflammation, sympathetic nerve hyperactivity, and metabolic disease are redundantly linked. In keeping with the theory that hypothalamic dysregulation is a primary event driving METS, exogenous induction of hypothalamic inflammation recapitulates many features of METS, whereas CNS-penetrant anti-inflammatory agents ameliorate many features of METS[5, 6]. Centrally acting sympatholytic agents comparably improve hypertension, hyperlipidemia, and insulin resistance[7, 8]. Strikingly, many studies have induced central inflammation using dietary or pharmacologic approaches. However, comparatively few studies have explored the benefits of enhancing endogenous regulatory mechanisms, especially those which appear to be damaged by the underlying disease process. Of these potential mechanisms, the somatostatinergic (SST) system is a particularly attractive therapeutic target because it both negatively regulates sympathetic outflow[9–16] at many levels of the neural axis and frequently inhibits inflammatory cytokine secretion. Most critically, hypothalamic SST neurons are depleted in preclinical models of METS/Type 2DM [17]. These findings suggest dysfunction of SST neurons could be an aggravating factor in METS hypothalamic inflammation and sympathetic nerve hyperactivity, a theory supported by the finding that SST suppresses cytokine release in multiple tissues [18, 19]. Exogenous SST and its analogues reduce peripheral sensory nerve hyperactivation in the face of inflammatory injury[20, 21], and exogenous somatostatin receptor activation preserves the blood brain barrier against inflammatory damage[22]. Interestingly, current data on SST regulation of microglial and astrocyte cytokine release is conflicting and largely derived from cell culture studies, highlighting the need for in-vivo studies to clarify its effects on hypothalamic inflammation [23–25].

Because the hypothalamus is a master regulator of homeostasis, hypothalamic inflammation may induce profound dysfunction across many organ systems. Perhaps nowhere is this more concerning than in hypothalamic regulation of the immune system[26], because impaired regulation of peripheral immunity has the capacity to initiate positive feedback loops sustaining hypothalamic inflammation. Specifically, altered sympathetic outflow to lymphoid organs may increase granulocyte and monocyte precursor (GMP) production, driving myeloidosis observed in individuals with Type 1 and Type 2 diabetes, as well as animal models of Type 1 DM and METS [27, 28]. These immune cells may in turn infiltrate the hypothalamus, further dysregulating autonomic outflow, as has recently been shown in neurogenic hypertension[29].

Thus, hypothalamic dysfunction has the capacity to induce downstream injury directly through hyperactivation of sympathetic nerves, as well as indirectly, through the altering of immune homeostasis and initiation of systemic inflammation. For example, bone marrow neuropathy induced alterations in circulating immune cells, with increased pro-inflammatory and decreased vasoreparative cells, has been shown to be upstream of diabetic retinopathy in several preclinical models[17].

Other disruptions in neural regulation may result from direct nerve hyperactivity. For example, in several dietary models of METS, norepinephrine turnover (NETO) in adipose tissue is rapidly increased[30]. As few as three weeks of cafeteria diet feeding doubles the rate of norepinephrine turnover onto retroperitoneal fat and increases NETO by 60% in epididymal fat[31]. However, comparatively little data examines the consequences of this neurometabolic disruption on signature features of adipose tissue physiology, especially lipolysis. This gap is striking, insofar as much data indicate that the neural regulation of lipolysis is greatly disrupted in METS in both humans and animal models[32–37].

In this study, we proposed that loss of the endogenous hypothalamic inhibitory neurotransmitter and anti-inflammatory agent, SST, could recapitulate features of diabetic pathologies in the absence of hyperglycemia. Thus, through a reductionist approach of selective hypothalamic SST loss, we asked whether loss of this small but critical population of cells could recapitulate aspects of metabolic disease.

## Results

### Aged db/db Hypothalamus Exhibits a Loss of Somatostatinergic Neurons

Previous data indicated a reduction in hypothalamic somatostatin in BBZ/WoR rat hypothalamus[17]. We tested the hypothesis that this phenomenon was a general finding in METS/type 2 DM models by examining somatostatin cell density in the leptin receptor deficient db/db mouse model of METS and mild type 2 diabetes. Serial coronal sections of the entire paraventricular and periventricular nucleus were examined using immunohistochemistry for somatostatin (Figure 1a). Sections were oriented with reference to the Allen Brain Atlas. SST neuron density varied systematically between the paraventricular and periventricular nucleus, with an overall higher abundance of SST neurons in the periventricular nucleus (Figure 1B). A strong rostral-caudal gradient was observed in the periventricular nucleus, with the highest densities of SST neurons between Bregma coordinates −0.28 to −1.055 in db/m control mice hypothalamus. A significant reduction of SST neurons was observed at most Bregma levels in the periventricular nuclei of db/db mice compared to db/m control mice, whereas paraventricular nucleus SST density exhibited reductions at only one Bregma level.

**Figure 1.**
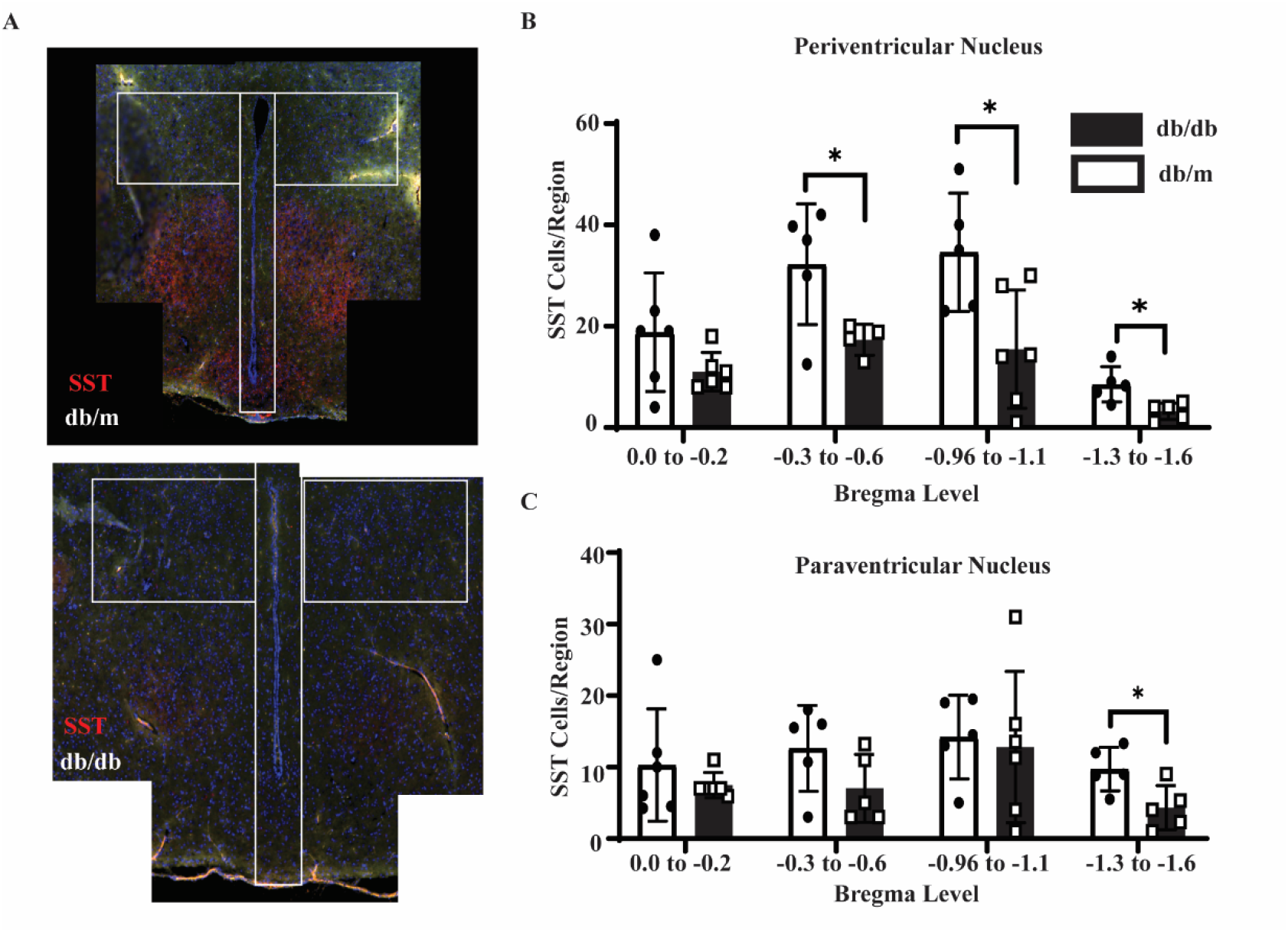
Somatostatin immunofluorescence across the entire hypothalamus paraventricular nucleus. db/db hypothalamus exhibits a reduction in immunofluorescent somatostatin staining at 12 months of age, as compared to heterozygote littermate controls (A). Cell counts of somatostatin positive nuclei in the periventricular nucleus from Bregma 0.0 to Bregma -1.6 (B). Cell counts of somatostatin positive nuclei in the paraventricular nucleus from Bregma 0.0 to Bregma -1.6 (C).

### AAV-DTA Treatment Depletes Somatostatinergic Cells and Induces Hypothalamic Inflammation

To mimic the observed reductions in somatostatinergic cells within the diabetic hypothalamus, SST-^cre^ mice received stereotactic injections of adeno-associated virus (AAV) containing a cre-driven cytotoxic alpha subunit of diphtheria toxin into the hypothalamus (SST-DTA). Control mice received a virus containing a GFP construct (SST-GFP/control). At 1-week post-treatment, a robust reduction of SST neurons was observed in the hypothalamus, but not in neighboring brain regions such as the ventromedial thalamus (Figure 2A). At 8 months post-DTA exposure, SST neuronal density remains suppressed (Figure 2B, Figure 2C). A smaller reduction of overall NeuN positive cells was observed (Figure 2D), indicating that SST expressing neurons were preferentially eliminated by this approach. This finding is strengthened by the observation that relativizing somatostatin density to overall NeuN density demonstrates a significant loss in SST-DTA mice (Figure 2E).

**Figure 2.**
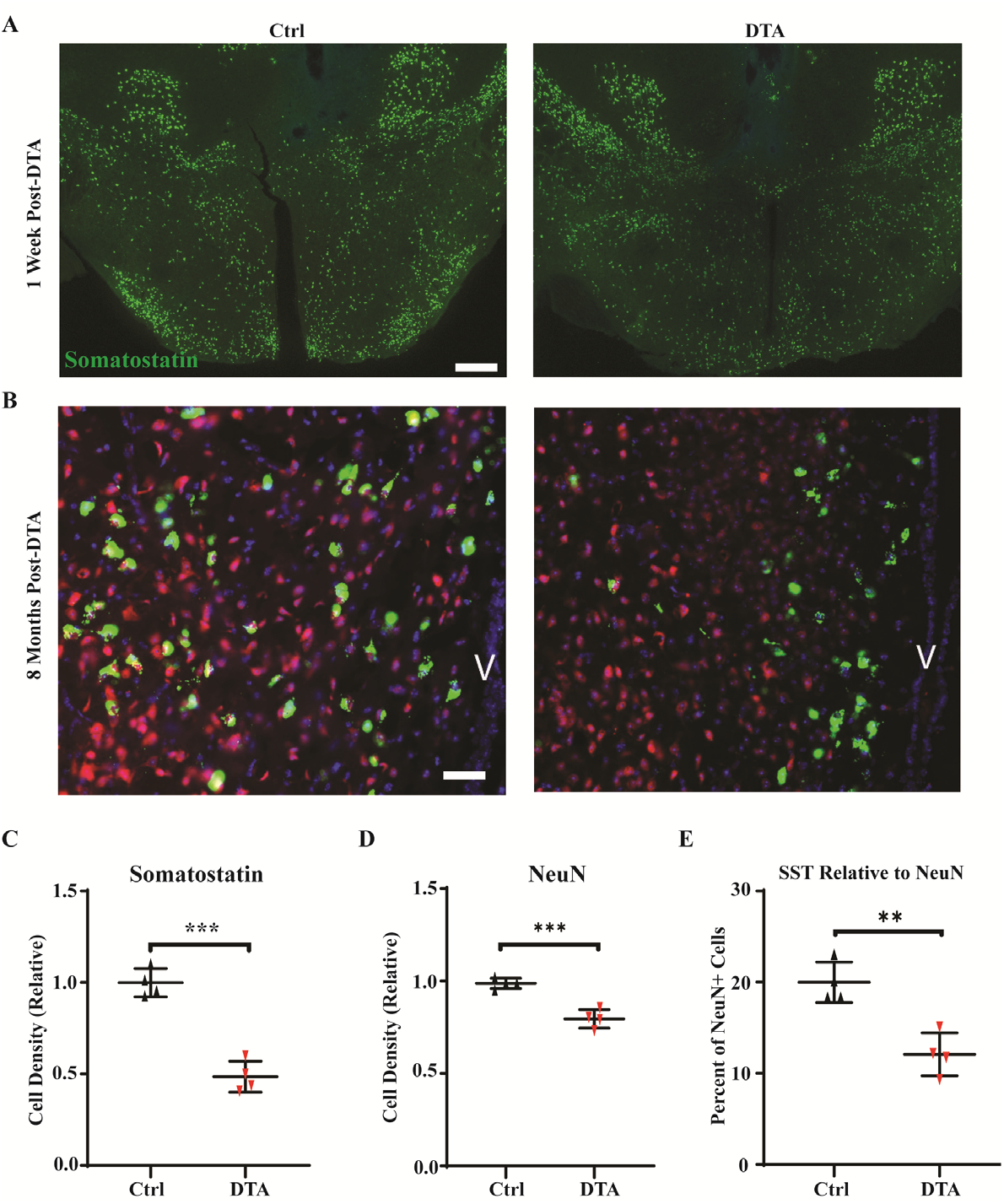
AAV-DTA delivery reduces hypothalamic somatostatinergic neurons. 1 week post-injection, a large reduction in somatostatin associated GFP is observed in hypothalamus, but not adjacent brain regions (A). Higher magnification images (40x) taken at the end of the study show sustained reductions in somatostatin associated fluorescence (green) in DTA treated mice, with small changes in NeuN positive neurons in red (B). Quantification of reduction in somatostatin in DTA treated mice, as compared to AAV-scrambleGFP mice; cell densities were calculated per field and relativized to control mice (C). Quantification of reduction in NeuN in DTA treated mice (D).

Concomitant with these findings, at 8 months DTA-treated mice exhibited an increased density of Iba1^+^ cells (Figure 3A). Iba1^+^ cells reflect microglial activation, with cells exhibiting more cytoplasm around the nucleus and short cell processes with loss of delicate dendritic ramifications compared to Iba1^+^ cells in controls. This increase in number and shape has been observed under disparate causes of central nervous system inflammation[38]. Additionally, SST-DTA mice exhibited signs of astrocytosis, measured via glial fibrillary acidic protein (GFAP), with qualitatively more complex branching patterns as well as quantitatively increased cell density (Figure 3B), reminiscent of findings in type 1 diabetes models, which also show elevations in hypothalamic Iba1^+^ cells [27]. Taken together, these data suggest that depleting hypothalamic SST-cells is sufficient to induce hypothalamic inflammation, and that this inflammation persists long after the initial DTA-mediated cell death.

**Figure 3.**
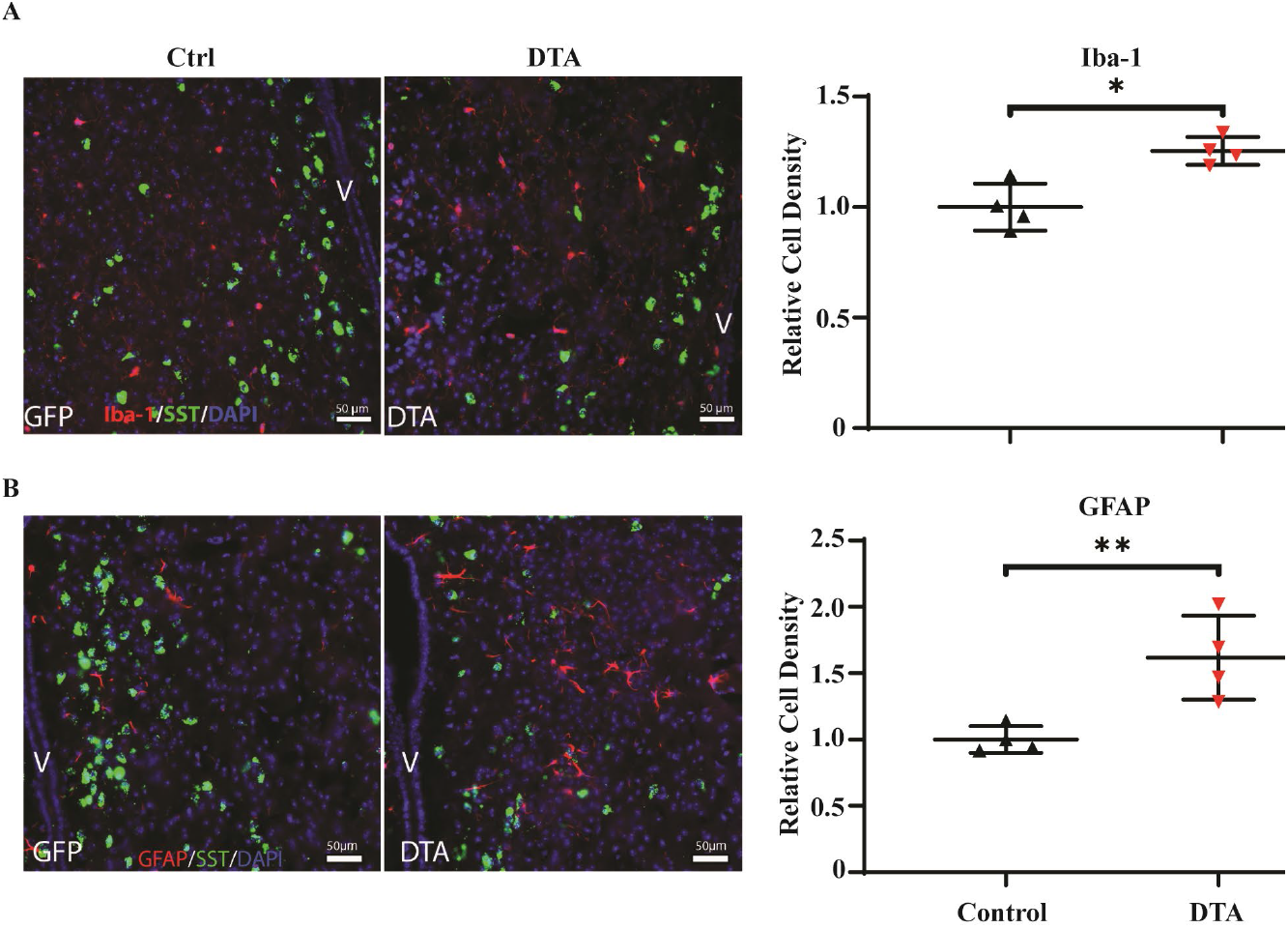
SST-DTA mice exhibit hypothalamic inflammation. Immunofluorescent images of Iba+ cells (red), somatostatin (GFP), and nuclei (DAPI). Cell densities were calculated per microscopic field and relativized to control mice. (A). Immunofluorescent images of glial fibrillary acidic protein (GFAP)+ cells (red), somatostatin (GFP), and nuclei (DAPI). Cell densities were calculated per microscopic field and relativized to control mice. (B).

A specific reduction of SST neurons is predicted to leave an enriched population of cells mediating sympathetic outflow without endogenous inhibitors. To test this hypothesis, immunohistochemistry for c-fos positivity in hypothalamic sections was undertaken (Figure 4A). Increases in c-fos staining are a commonly utilized proxy for recent cell activation[39]. Computing the ratio of c-fos positive cells in DTA mice versus controlled treated mice demonstrates a significant increase in the prevalence of c-fos positive neurons in SST-DTA mice (Figure 4B).

**Figure 4.**
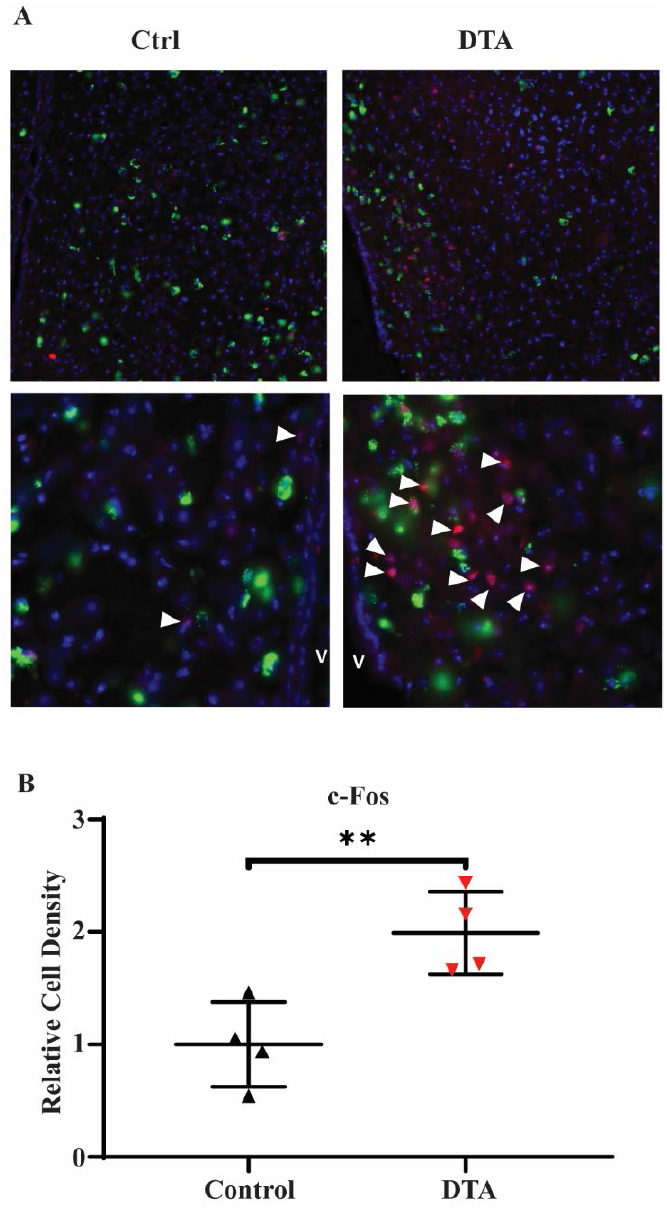
SST-DTA mice exhibit increased in periventricular hypothalamic c-Fos labeling. Representative micrographs at low power (5x) and higher power (20x) showing somatostatin associated GFP fluorescence and c-Fos labeling (red) in both DTA treated, and control treated mice (A). Quantification of relative cell density of c-Fos positive cells in DTA treated mice versus controls (B).

### SST-DTA mice recapitulates key diabetic metabolic phenotypes

**A**dipose tissue is broadly divided into brown and white adipose tissue, the latter of which is further subdivided into visceral and subcutaneous fat. Visceral fat is defined by venous drainage to the liver; in rodents, the only fat pad which drains to the liver is the mesentery[40]. This anatomical feature plays a key role in the development of insulin resistance[41, 42]. Adipose tissue releases stored fatty acids and glycerol in response to sympathetic nerve input, which is dependent on the hypothalamus[4, 35, 43]. Adipose tissue in individuals with METS and Type 2 DM, as well as preclinical models, has a reduced lipolytic response to neural input [36, 37]. To measure neurally driven lipolysis, we established a dose-response curve of acute adipose tissue explants in response to isoproterenol, a β-AR agonist, finding that adipose tissue from mesenteric or inguinal subcutaneous origin released free fatty acids and glycerol dose-responsively to isoproterenol stimulation. This approach assays the sensitivity of adipose tissue lipolysis to catecholamines (Figure 1A). Using this method, we measured catecholamine sensitivity in db/db mice. Compared to their heterozygote and littermate controls, 6 to 12 month old db/db mice exhibited catecholamine resistance in both mesenteric and subcutaneous fat (Figure 5B). We then compared the lipolytic response of adipose tissue from DTA and control treated animals. Interestingly, inguinal subcutaneous fat from DTA-treated animals exhibited similar catecholamine sensitivity to GFP virus-treated control animals. In contrast, catecholamine induced lipolysis was suppressed by 25% in the mesenteric fat of SST-DTA mice (Figure 5C). Because chronic local release of NE could in theory drive desensitization to catecholamines, we examined the NE concentration in inguinal subcutaneous fat and mesenteric fat, finding that mesenteric fat trended towards higher levels of NE than subcutaneous fat (Figure 5D).

**Figure 5.**
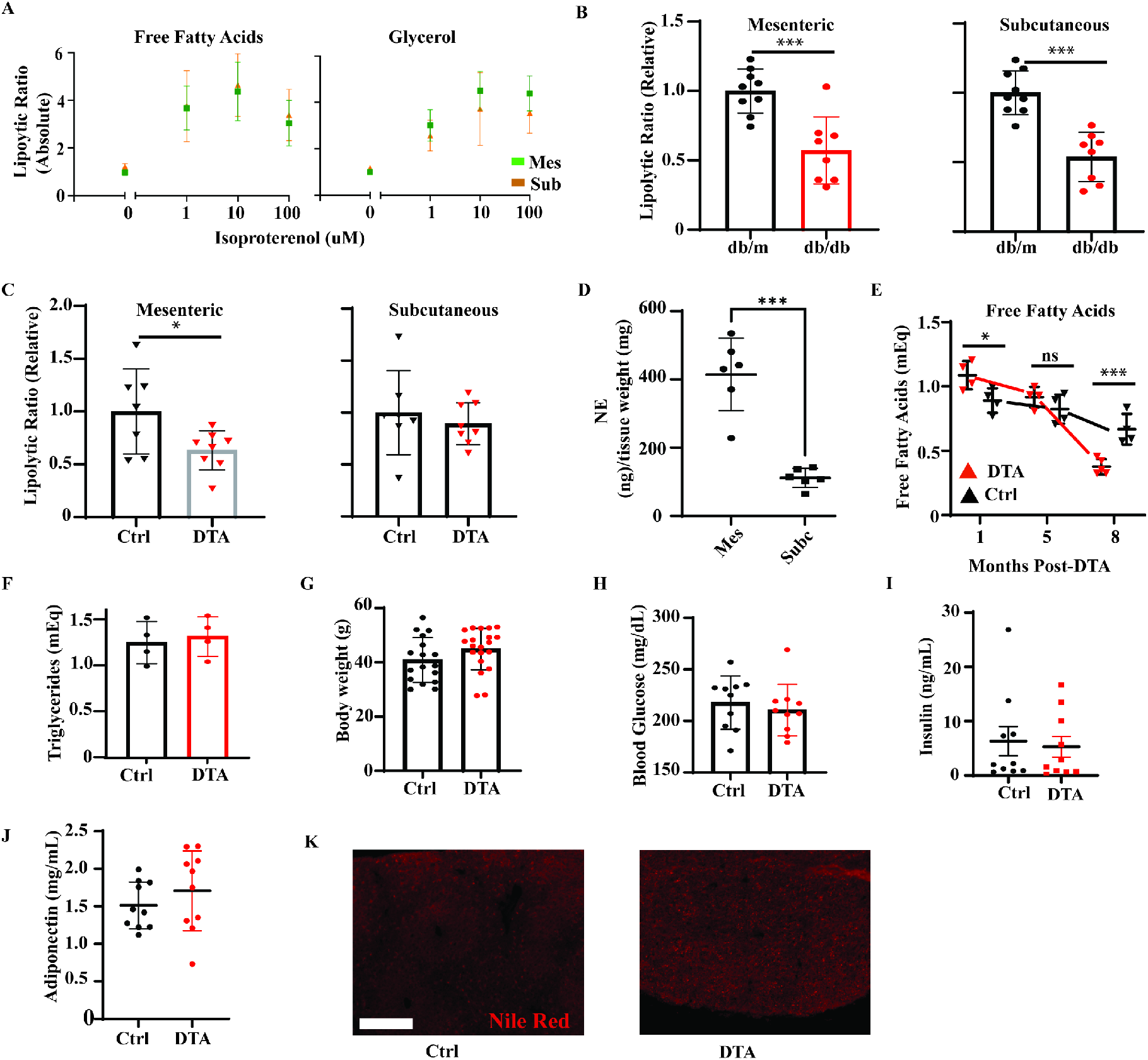
SST-DTA mice exhibit key diabetic metabolic phenotypes. Isoproterenol dose-dependently drives the release of free fatty acids and glycerol from acute fat explants (A). Plots of ex-vivo lipolysis in mesenteric and inguinal subcutaneous fat from 12 month old db/db mice. Data are relativized to heterozygote control littermates (B). Plots of ex-vivo lipolysis in mesenteric and inguinal subcutaneous fat at 8 months post-DTA (C). Norepinephrine concentration in wild-type mesenteric and subcutaneous fat measured by HPLC and electrochemical detection (D). Serial plasma free fatty acid measurements in DTA and control treated mice show dynamic changes dependent on duration of hypothalamic injury. (E). Triglyceride measurements in control and DTA treated mice show no changes at 8 months post-injection (F). Body weight measurements at 8 months of age show no overall trend (G). Random blood glucose measurements at 8 months of age show no effect of DTA treatment (H). Plasma insulin under the fed state is not elevated in DTA mice (I). Plasma adiponectin is not depleted in DTA mice (J). Nile Red neutral lipid staining in DTA and control livers does not show gross lipid deposits; scale bar is 50 microns (K).

Adipose tissue lipolysis releases free fatty acids (FFA) and glycerol into the circulation and is largely controlled by sympathetic nervous system input [43]. Thus, reduced plasma free fatty acids are a plausible correlate of blunted responses to sympathetic nerve input. Indeed, at 8 months of age, SST-DTA mice exhibited a 40% reduction in plasma free fatty acids (Figure 5E). Examining FFA content of archived plasma samples from monthly blood draws showed a striking pattern of early elevations in FFA content at 1 month post-DTA, followed by comparable FFA levels at 5 months post-DTA, and culminating in the aforementioned reduction at 8 months (Figure 5E).

Elevations of FFAs in the plasma of METS and type 2 diabetic individuals are driven by increased *basal* lipolysis, so much so that even non-diabetic individuals with a family histories of diabetes exhibit enhanced basal lipolysis[32]. Indeed, both insulin resistance and enhanced basal lipolysis are driven by elevations in TNF-a [44]. In contrast, we observe a deficit in *induced* lipolysis. Clinically, lipolytic function correlates with other plasma lipids (triglycerides and HDL;[34]). In our hands, however, we observed no differences in triglycerides between SST-DTA mice and control treated animals (Figure 5F).

Notably, no trends in weight gain or loss were observed in SST-DTA mice (Figure 5G), nor were any differences observed in random blood glucose measurements (Figure 5H) or insulin levels (Figure 5I). Resistance to catecholamine induced lipolysis has been linked with a loss of adipose adiponectin secretion. This loss is an important feature of diabetes and cardiovascular disease, as adiponectin is an anti-inflammatory, insulin sensitizing hormone[45–47]. However, no differences in circulating adiponectin levels were observed in SST-DTA mice (Figure 5J), though we did not measure the concentration of adiponectin in the hepatic portal vein, to which mesenteric fat drains. Free fatty acids released by adipose tissue are cleared by the liver; as such, we assay liver neutral lipids using Nile Red neutral lipid staining. No gross lipid deposits were observed in SST-DTA liver (Figure 5K).

### SST-DTA Mice Exhibit Increased Granulocyte Precursors in Bone Marrow and Increased Inflammatory State of Bone Marrow and Circulating Monocyte Pool

In preclinical models of diabetes, hematopoiesis is biased towards myeloid progenitors with subsequent alterations in peripheral blood monocytes [48–51]. Indeed, numerous changes have been observed in diabetic hematopoiesis, even at very early stages. Specifically, increased short term repopulating hematopoietic stem cells and suppressed long term repopulating hematopoietic stem cells were observed, which may further bias hematopoiesis towards myeloid lineages[48, 52]. Flow cytometry of lineage negative, sca1^-^, c-kit^+^ cells from SST-DTA and control mice showed an increased percentage (relative to all lin^-^, sca1^-^, c-kit^+^ cells) of granulocyte monocyte precursors (GMP) cells (CD34^+^, CD16/32^+^) (Figure 6A,B). In contrast, a decreased percentage of megakaryocyte erythrocyte precursors (MEP) was observed in SST-DTA mice (CD34^-^, CD16/32^-^) (Figure 6C). Relative monocyte increases (relative to total CD45^+^ cells) were observed (Figure 6D,E). Additionally, SST-DTA mice exhibited increased percentages of proinflammatory CCR2^+^ monocytes (Figure 6F). Examining monocyte subsets by Ly6c expression revealed an expansion (relative to all CD45^+^ cells) of the Ly6c^+^ population in SST-DTA mice, as compared to GFP treated controls.

### SST-DTA Mice Exhibit Increased Retinal Inflammation and Reduced Retinal Function

Previously, we reported that two different dysfunctions in the neural axis, hypothalamic inflammation and bone marrow neuropathy, precede retinal microvascular disease and reduced retinal function[6, 27]. To assess whether hypothalamic SST-depletion recapitulated these findings, visual function and flow cytometry of the neural retina were performed at 8 months post-treatment (9 months of age). SST-DTA mice exhibited higher levels of retinal monocytes than control mice (Figure 7A).

**Figure 7.**
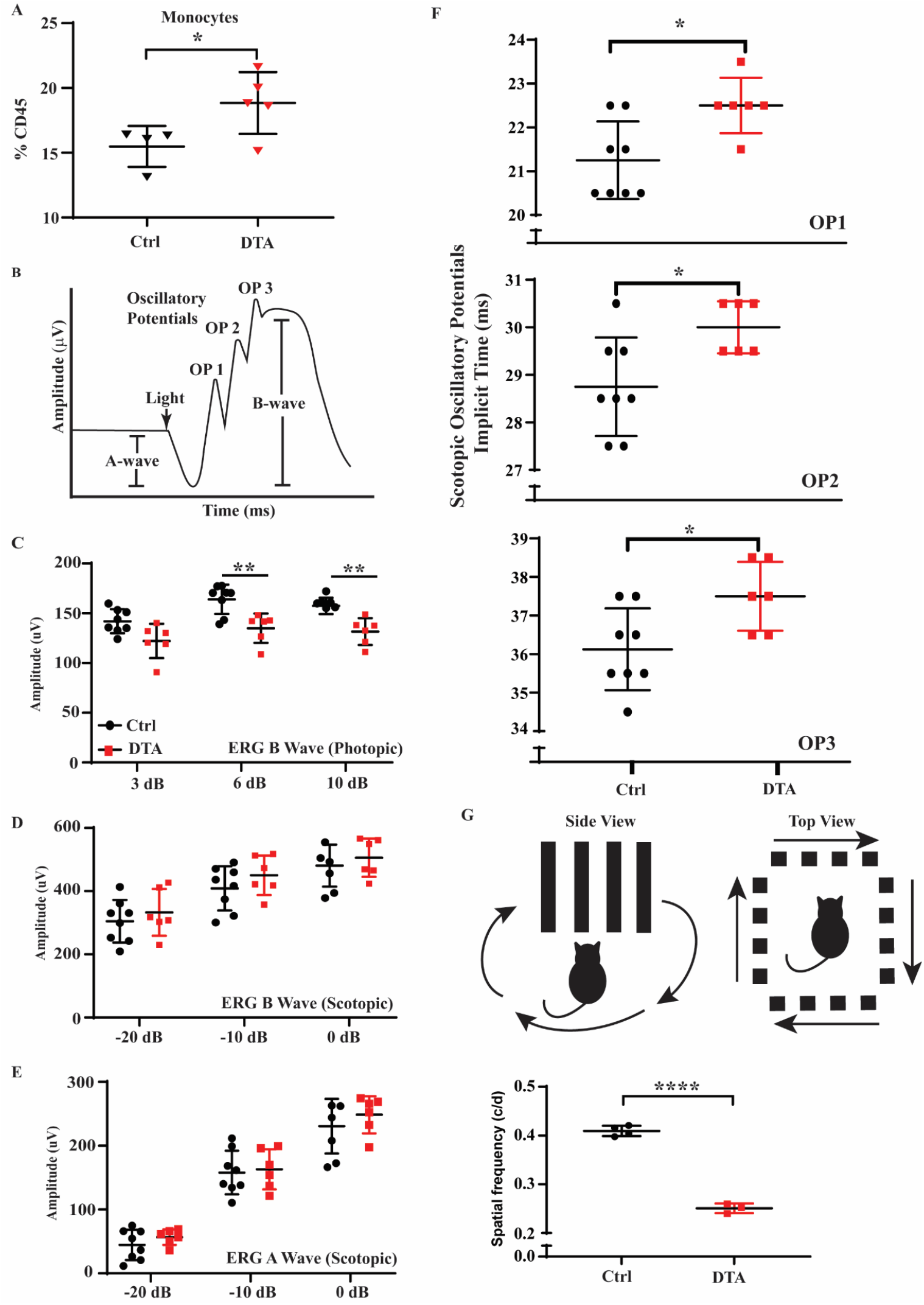
SST-DTA mice exhibit changes in retinal immunology and function. Results from flow cytometry show increases in retinal monocytes in DTA treated mice (A). Schematic diagram of typical ERG response (B). Electroretinogram (ERG) recordings from DTA mice exhibit deficits in B-wave responses under photopic conditions (C), but not scotopic conditions (D). A-wave abnormalities are not observed (E). DTA mice show delayed oscillatory potential implicit times at the first, second, and third oscillatory potentials (F). Schematized optokinetic diagram and data show that the minimum detectable cycles per degree, a measure of spatial resolution, are lower in DTA treated mice (G).

Simultaneously, these mice exhibited dysfunctional electroretinographic (ERG) responses. The electroretinogram assays the electrical response of the retina to light (Figure 7B). The ERG is summed field potential containing a photoreceptor driven A-wave (driven by a hyperpolarizing response to light) and a neural retina (retinal bipolar cell) dominant B-wave, driven by a depolarizing response to photoreceptor synaptic input. Scotopic ERG’s are conducted using low light intensities and largely reflect rod photoreceptor activity. In contrast, photopic ERGs isolate cone photoreceptor activity by delivering a chronic background light to suppress rod photoreceptor activity. Cone photoreceptors are then stimulated by superimposing a bright light. In addition to A and B waves, ERGs exhibit oscillatory potentials (OP); OP have complex, incompletely described origins in the neural retina. Importantly, disruptions in the timing of OP is an early event in diabetic retinal dysfunction which occurs prior to frank retinopathy. SST-DTA mice exhibit disruptions in photopic B-waves (Figure 7C), but not photopic A-waves or scotopic B-waves (Figure 7D,E). Delays in the onset (implicit) time of each oscillatory potential were detected in SST-DTA mice (Figure 7F). Finally, in a second measure of retinal function, we examined optokinetic tracking, a behavioral measure of visual acuity. SST-DTA mice exhibited profound reductions in spatial acuity (Figure 7G) from 0.4 cycles/degree in control mice to 0.25 cycles per degree in SST-DTA mice, p>0.0001).

## Discussion

The main findings of our study support the hypothesis that SST neurons are damaged in murine models of METS and type 2 diabetes (Figure 8). The loss of hypothalamic somatostatinergic cells has been observed in T2D rodent models and has been implicated in diabetic complications pathogenesis, in particular development of microvascular complications such as diabetic retinopathy[17]. We observed a robust reduction in hypothalamic SST neurons within 1 week of stereotactic injections of an AAV containing a floxed DTA toxin construct to SST^cre^ mice. These reductions in hypothalamic SST neurons were sustained at 8 months post-DTA. SST neural ablation recapitulated several pathologies observed in diabetes and METS. SST-DTA mice exhibit neuroinflammation comparable to that observed in diabetes, as demonstrated by increases in Iba-1^+^cells[6], astrocyte activation, and increases in c-fos expression.

**Figure 8.**
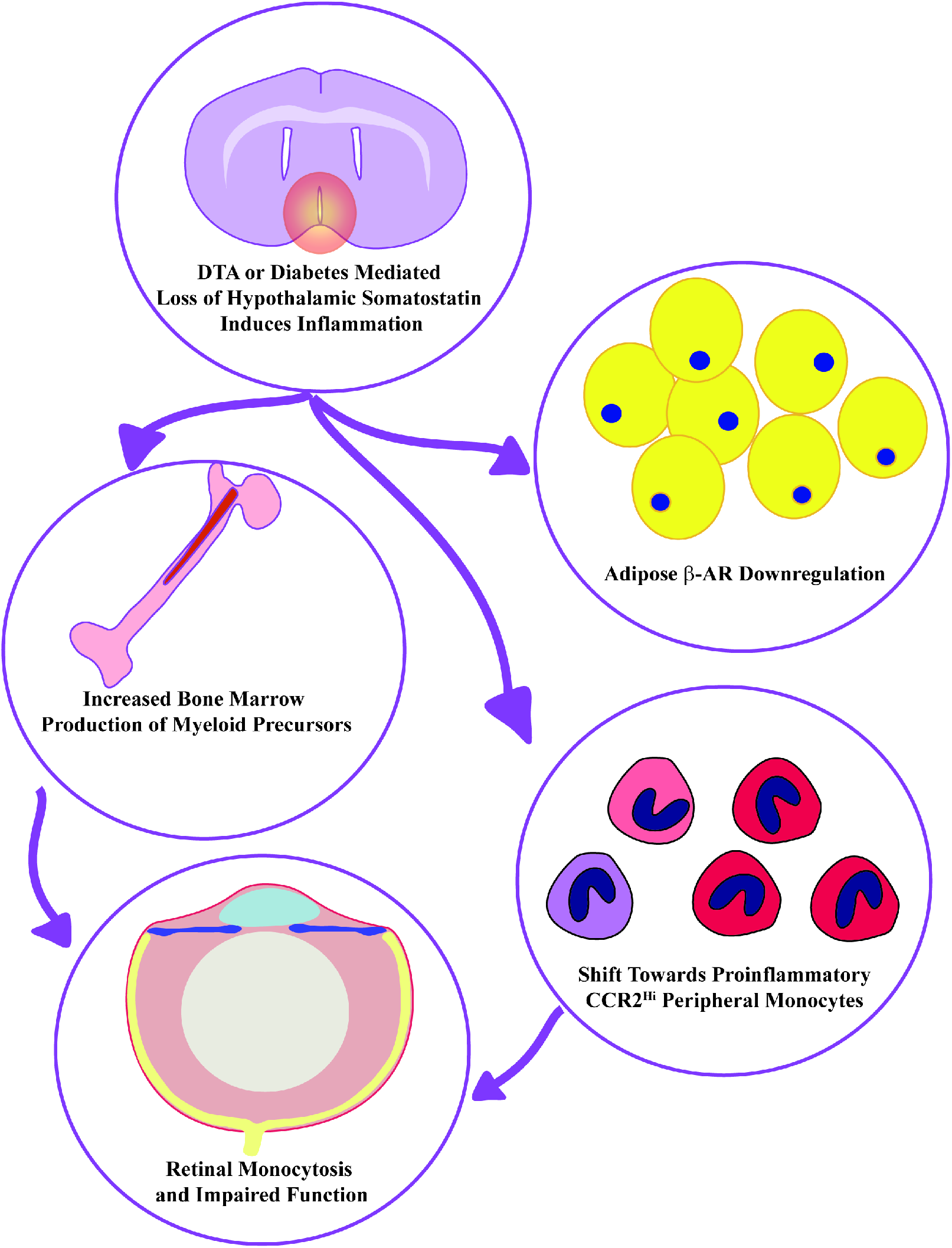
Schematic showing downstream impacts of diabetes or DTA mediated loss of hypothalamic somatostatinergic neurons. These deficits include altered hematopoiesis, increases in CCR2+ proinflammatory monocytes, loss of retinal function with concomitant retinal monocytosis, as well as resistance to neural control of lipolysis in mesenteric adipose.

The loss of hypothalamic SST neurons resulted in marked changes in hematopoiesis, specifically, a myeloidosis of the bone marrow and alterations in circulating blood monocytes. We observed increases in retinal monocytes in SST-DTA mice, comparable to infiltration observed in diabetic models [53, 54]. SST-DTA mice also exhibited impaired catecholamine sensitivity in their mesenteric adipose. Catecholamine resistance is a common finding in mouse models and patients with METS and T2D[37, 55, 56]. While all these features are found in diabetes, the SST-DTA mice exhibited normal glucose homeostasis. These findings support the theory that disruptions in neural regulation of immunity and metabolism are early features of metabolic disease. For example, previous data indicated that bone marrow neuropathy precedes retinopathy, the most common microvascular injury in diabetes[57]. Alterations in bone marrow innervation were theorized to alter the balance of the release of vasoreparative and pro-inflammatory cells into the circulation.

The present data link these later stage observations to early, upstream hypothalamic injury, which largely recapitulate the immunologic findings made in mouse models of diabetes exhibiting bone marrow neuropathy. These findings make sense in light of the unique time course of pathology observed all levels of the neural axis. For example, mice subjected to high fat diet (HFD) show hypothalamic inflammation after 24 hours, particularly in astrocytes[3]. Inflammation is closely followed by brain insulin resistance [58]. After months of HFD, the hypothalamus exhibits apoptotic cell death [59] and reduced neurogenesis[60]. Genetic models of METS recapitulate the sequence of hypothalamic inflammation, insulin resistance [61] and cell death [62]. Progressive dysfunction in the hypothalamus mirrors alterations in the peripheral nervous system. For example, peripheral nerve hyperactivity may progress to the bone marrow neuropathy observed in both streptozotocin (STZ)-treated mice and BBZ/WoR type 2 diabetic rats [27, 57]. Notably, the bone marrow-brain relationship is bidirectional, as ablation of bone marrow beta-adrenergic receptors induces hypothalamic inflammation, emphasizing the reciprocal feedback loops linking central nervous system inflammation and peripheral immunity[63].

The present data inform our understanding of the physiology of SST neurons in the brain. While somatostatin uniformly suppresses cytokine release in the periphery, anti-inflammatory action in the brain is more controversial and largely derived from cell culture studies[20, 23-25]. The data presented here suggest that somatostatinergic neurons, whether through SST itself or other mechanisms, repress CNS inflammation. This constitutes an important functional contribution to the literature, which has thus far not definitely resolved somatostatin’s role in the regulation central nervous system inflammation. In addition to their anti-inflammatory function, SST neurons also secrete the classically inhibitory neurotransmitters GABA and glycine in brainstem and cortex [9], though recent data indicate that some hypothalamic SST neurons are glutamatergic [64], whereas a majority of spinal SST^+^ cells are glutamatergic[65]. In brief, the molecular heterogeneity of SST neurons requires functional characterization.

We show that depletion of hypothalamic SST neurons increases local c-Fos expression, potentially suggesting release of presympathetic hypothalamic nuclei from inhibition. In both METS and type 2 diabetes, elevations in sympathetic nervous system activity are observed[66].

Thus, recapitulating pathologies observed in these diseases would support the hypothesis that hypothalamic SST neurons are an endogenous negative regulator of SNS outflow lost in diabetes.

Indeed, we observe a well-characterized hallmark of type 2 diabetes and METS in the adipose tissue of SST-DTA mice, catecholamine resistance. Reduced lipolytic response to β-adrenergic stimulation has been measured in diabetic individuals and in preclinical models[37, 55, 56]. In-vitro approaches have shown that incubation of adipocytes with TNF-α is sufficient to reduce responses to β-adrenergic agonists[67], suggesting that chronic low-grade inflammation could induce or aggravate catecholamine resistance. However, receptor down-regulation in response to excess ligand is common, and under dietary challenge, adipose tissue sympathetic nerves release excess norepinephrine[30, 68]. Thus, hyperactivity of adipose sympathetic nerves (and resultant catecholamine excess) is a plausible mechanism for catecholamine resistance. In DTA-treated animals, we observe a strong reduction of catecholamine induced lipolysis comparable to that observed in METS and type 2 diabetes. Surprisingly, catecholamine resistance was restricted to the mesenteric adipose. Under caloric restriction, visceral adipose is a preferential site of weight loss with high levels of norepinephrine turnover [69]. Mesenteric fat is also richly represented at the level of the hypothalamic PVN, the primary site of our manipulation[40]. We speculate that richly innervated mesenteric adipose is uniquely vulnerable to desensitization by sympathetic hyperactivation, and that this underlies the fat depot specific effect observed here. In support of this view, we assessed local concentrations of norepinephrine in mesenteric fat and subcutaneous inguinal fat, finding elevated concentrations of NE in mesenteric fat.

Several lines of data indicate that diabetes disrupts bone marrow innervation. Models of type 1 and type 2 diabetes show decreased bone marrow tyrosine hydroxylase and CGRP [27, 57]. Loss of hypothalamic somatostatinergic neurons is theorized to hyperactivate the bone marrow circuit, contributing to this neuropathy[17]. Alterations in bone marrow stromal cells and in hematopoiesis have been attributed to this neuropathy, in particular as an upstream driver of diabetic monocytosis[6, 27]. In this study, depleting somatostatinergic cells from the hypothalamus recapitulated many of these findings. Flow cytometry studies of the bone marrow demonstrated a hematopoietic bias towards monocytes and granulocyte precursors, recapitulating findings recently published in db/db mice[70]. Peripheral blood monocytes were preferentially proinflammatory and CCR2^hi^, with an expansion of monocytes (relative to CD45^+^ cells) as well as increase in Ly6C^hi^ monocytes.

Retinal dysfunction, mediated by vascular injury and immune infiltrate, is a common feature of diabetes. We observe increases in retinal monocytes in SST-DTA mice, comparable to infiltration observed in diabetic models. This finding was accompanied by reductions in spatial acuity, by optokinetic tracking, and neural retinal function which were comparable to earliest changes identified in the diabetic retina, which directly precede frank retinopathy. For example, we detected delays in the implicit times of ERG oscillatory potentials, which is the most common ERG dysfunction identified in diabetic patients[53]. In animal models, delayed OP implicit time precedes retinal vascular damage in chronically high fat diet fed mice[54]. Notably, these mice exhibited a pro-inflammatory retinal phenotype prior to the onset of ERG dysfunction, suggesting a step-wise pathophysiology linking inflammation to retinal dysfunction and eventual vascular damage.

In summary, a reductionist approach of somatostatinergic cell deletion from the hypothalamus was capable of recapitulating many diabetic phenotypes. The significance of this finding is that this small but significant population of neurons impacts many target tissues of diabetic complications including, as we have shown here, the retina, brain, bone marrow, and adipose tissue. Pathology resulting from the loss of SST can be potentially corrected in diabetes by pharmacological strategies, for example, the use of a somatostatin analogues such as Octreotide, delivered intranasally, to directly target SST loss in the hypothalamus[71].

## Methods

### 1. Mice

All protocols were approved by the University of Alabama at Birmingham (UAB) IACUC. SST reporter mice were bred on a C57Bl6 background by Jax laboratories in their animal facility under a 12:12 light dark cycle by crossing Sst^tm2.1(cre)Zjh^ (Jax 013044) to the cre-dependent reporter line B6.Cg-Gt(ROSA)26Sor^tm6(CAG-ZsGreen1)Hze^ (Jax 007906). Only mice which were heterozygote at both genomic insertions were used. Stereotactic injections (100-200 nL per side) were performed by a trained operator at the University of Alabama at Birmingham. All animals were given presurgical analgesics and anesthetized with isoflurane. The following stereotactic coordinates were used: (Anteroposterior: −0.5; Mediolateral: +/− 0.220; Dorsoventral: −4.8) in mice aged 8 to 10 weeks. The control AAV virus (AAV-GFP) contained the following genetic construct: an hSyn promoter, GFP flanked by inverted LoxP sites, and a 5’ WHP post-transcriptional regulatory element (WPRE). Control mice were defined as SST-reporters injected with the control virus, as outlined above. The DTA virus (AAV-DTA) contained a CMV promoter and a floxed mCherry stop codon upstream of the coding sequence for the cytoxic subunit A of the diptheria toxin, and a WPRE. The virus was a gift from Patrick M. Fuller [72, 73]. Animals were group housed and fed chow diet ad libitum. Male mice were used for immunologic studies and mice of both sexes were used for metabolism studies.

### 2. Immunohistochemistry

Mice were euthanized using isoflurane and cervical dislocation. Brains were dissected into PBS, fixed in 4% paraformaldehyde for 24 hours at 4°C and transferred to 30% sucrose for 24 hours. Brains were embedded in optimal cutting temperature (OCT) media and sectioned at 14 uM thickness and stored at −20°C until labeling. Brain sections were washed twice in PBS for 10 minutes and blocked in 1% bovine serum albumin (BSA) in 0.3% Triton-X for 30 minutes at room temperature. The following primary antibodies were incubated overnight: rat anti-somatostatin (M09204, Abcam,1:100), rabbit anti-NeuN (PA5-78499, Thermo Fisher Scientific; 1: 100), rabbit anti-c-Fos (ab90289, abcam, 1:100), rabbit anti-Iba-1 (019-19741, Wako, Osaka, Japan, 1: 200), and rabbit anti-GFAP (Rb, ab33922, Abcam, 1:100). Sections then transferred to species-specific secondary antibodies conjugated with either AlexaFluor 594 (1:200) or AlexaFluor488 (1:100, Invitrogen-Molecular Probes, Carlsbad, CA) for 1.5 hours in room temperature. Following two additional PBS washes, sections were reacted with DAPI (D3571, Invitrogen by Thermo Fisher Scientific, 1:10000) for 10 minutes and mounted in antifade mounting medium (Victory). Somatostatin visualization required an antigen retrieval process in citrate buffer (Sigma Aldrich C8532, pH 6.0) for 30 minutes prior to blocking. Liver Nile Red (12 ug/mL; Thermofisher N1142) staining was performed for two hours at room temperature. All sections were examined using fluorescence microscopy with a Zeiss Axio Imager Z2. The images were analyzed by using Zen lite software for colocalization analysis. Cell counting was performed in ImageJ.

### 3. Adipose Tissue Lipolysis

Mesenteric and subcutaneous inguinal fat pads were dissected into PBS, cut into 50 to 100 mg pieces, and loaded into 96 well plates with basal lipolysis media (low glucose DMEM with 4% Fatty Acid Free BSA). Tissue was incubated at 37 °C with 4% CO_2_ for between 1 and 3 hours in basal media, then transferred to a new 96 well plate containing induced lipolysis media for 1 hour. Induced lipolysis media was composed of low glucose DMEM with 4% Fatty Acid Free BSA, 10 uM Triacsin C to inhibit fatty acid re-esterification and 10 uM isoproterenol to stimulate β adrenergic receptors. At the termination of the experiment, tissue was snap frozen and media was stored at 4°C until assayed for free glycerol. Free glycerol was measured with the Sigma Free Glycerol Assay Kit (Sigma MAK 117); colorimetric detection was performed using a Biotek Epoch microplate spectrophotometer at 550 nM. Lipolysis was calculated as the ratio of free glycerol in the induced media to the free glycerol in the basal media, normalized for duration of basal lipolysis[33].

### 4. Fatty Acid Measurements

Plasma free fatty acid were measured using Wako Diagnostics Non-Esterified Free Fatty Acid Kit (999-34691), following kit directions exactly. Absorbance at 540 nm was measured using a Biotek Epoch microplate spectrophotometer.

### 5. Flow Cytometry

#### Peripheral blood flow cytometry

Blood was collected via cheek bleed into heparinized capillary tubes under isoflourane anesthesia on a monthly basis. Blood was centrifuged at 3000 RPM at 4°C for 7 minutes, plasma was collected, packed cells were resuspended in cold ammonium chloride potassium (ACK) lysis buffer and lysed for 15 minutes on ice, then centrifuged at 400g at 4°C for 5 minutes. Cells were washed twice as follows: supernatant was decanted, the cell pellet was resuspended in 5 ml of sterile, filtered BEP (2% Fatty Acid Free BSA, 1% EDTA, in PBS), and centrifuged at 400g for 5 minutes. Following washes, cells were stained with custom antibody cocktails (see Table 1 for flow cytometry panels) for 20 minutes at room temperature in the dark, then washed twice more with BEP as previously described. Flow cytometry was performed using a BD Celesta and BD FACSymphony. All antibodies are listed in Supplemental Table 1. Gating schema are shown in Supplemental Figure 1.

#### Bone marrow flow cytometry

Whole femurs were cleared of musculature and connective tissue using sterile gauze. A small portion of the distal and proximal epiphysis was resected, and the bone marrow was flushed using 6mL of sterile FEB buffer to generate a single cell suspension, then stored on ice until centrifuged. Red blood cell lysis was achieved with ACK lysis buffer for 15 minutes on ice and arrested with 10mL FEB buffer, before centrifugation and two washes with FEB prior to staining.

#### Retina flow cytometry

Retina was isolated and stored on ice until digestion with 1 mL RPMI containing 5% fetal bovine serum supplemented with Collagenase D and DNase. Retinas were incubated in this medium at 37°C for an hour and vortexed every fifteen minutes. The resulting suspension was passed through a 70 um cell straining with 10ml sterile FEB buffer, then centrifuged at 400g for 5 minutes at 4°C. The cell pellet was resuspended into 200 uL of FEB, stained, and flowed as outlined above.

### 6. Electroretinogram

Electroretinograms (ERGs) were recorded using the LKC Bigshot ERG apparatus. Mice were dark-adapted overnight, anesthetized with ketamine-xylazine in sterile 0.9% saline (80 mg/kg and 15 mg/kg total body mass, respectively) and then dilated with atropine/phenylephrine under dim red light. Contact lens electrodes were placed with Gonak. A steel reference electrode was placed subcutaneously on the dorsal aspect of the head, and a steel ground electrode was placed in the foot. Animals were exposed to 5 full-field white light flashes at 0.25 and 2.5 cd.s/m^2^ (scotopic condition), light-adapted for 5 min and exposed to 10–15 full-field white light flashes at 10 and 25 cd.s/m^2^ (photopic condition).

### 7. Optokinetic Tracking Responses

A trained observer-operator recorded the visual acuity of the mice as determined by observing their reflexive head tracking movements (optomotor response) while masked to the cohorts of the mice under examination. This was conducted using a virtual optomotor system (OptoMotry; Cerebral Mechanics, Inc., Lethbridge, Canada) using their custom software (OptoMotry HD software version 2.0.0(4907). The mice were placed on a platform inside a four walled chamber made of computer monitors with screens displaying vertical alternating black and white stripes (gratings), shown under photopic conditions at 100% contrast and with thicknesses (spatial frequencies) ranging from 0.2 to 0.5 cycles per degrees rotating at 12.0 d/s. In mice, optomotor behavioral assays measure the rotation of the head in response to the rotation of these gratings. Head rotation is driven by oculomotor reflex, which keeps moving images stable on the retina. The thickness of the gratings corresponds to spatial frequencies (cycles per degree). For example, thin gratings test high spatial frequencies and thick gratings test low spatial frequencies. The highest spatial frequency at which a head rotation could be reliably detected was recorded as the visual acuity of the mouse.

### 8. Statistical Analyses

All statistical analyses were conducted using GraphPad Prism 8.2.0. Two-sample t-tests were used to detect significant differences (greater than p=0.05) between SST-DTA and SST-GFP groups.

### 9. Data and Resource Availability

The data sets generated and/or analyzed during the current study are available from the corresponding author on reasonable request. No applicable resources were generated during the current study.

## Acknowledgments

This study was supported by the National Institutes of Health Grants R01EY028037 to M.B. Grant and T32 GM008361 Medical Scientist Training Program to William Geisler, and Research to Prevent Blindness-Unrestricted Grants. We gratefully acknowledge Patrick M. Fuller for the gift of the cre-inducible DTA toxin virus.

## Author Contributions

CH managed all experiments and data collection and edited the manuscript. RFR collected metabolic data and wrote the manuscript. RB performed the stereotactic surgery. PH performed the tissue processing and immunohistochemistry. YAA performed optokinetic and electroretinogram recordings, and CPV and ALFL assisted with flow cytometry and bone marrow function assays. GML, PMF, KLG aided in experimental design. MBG conceptualized experiment, obtained funding, designed experiments, assisted with data interpretation, edited the manuscript, and is the guarantor of all data in the manuscript.

## Competing Interests

The authors have no competing interests to announce with respect to the present paper.

---

**Supplementary Table 1:** Flow Cytometry Antibodies

| Bone Marrow-Lineage |  |  |
| --- | --- | --- |
| Target | Catalog Number | Fluorophore |
| c-Kit | Biolegend: 105841 | BV 785 |
| Lineage | Invitrogen 88-7772-72 | eFluor 450 |
| Ly6A/E (Sca1) | eBioscience:64-5981-82 | SB 645 |
| CD16/32 | eBioscience: 63-0161-82 | SB 600 |
| CD34 | BD: 560518 | APC-R700 |
| CD135(FLT3) | Invitrogen: 17-1357-41 | APC |
| Viability (FVD506) | eBioscience: 65 0866-14 | BV 510 |
| Bone Marrow-Monocytes |  |  |
| CD45 | eBioscience: 47045180 | APC-Cy7 |
| CD11b | BD: 563553 | BUV396 |
| Ly6G | BD: 741994 | BUV 805 |
| CD11c | BD: 563735 | BV786 |
| CCR2/CD192 | BD 747964 | BV711 |
| Retina-Monocytes |  |  |
| F4/80 | eBioscience: 48-4801-82 | eFluor450 |
| CD45 | eBioscience: 64-0451-82 | BV650 |
| CD11b | eBioscience:78-0112-82 | BV786 |
| Ly6G | Biolegend: 127639 | BV605 |

**Supplemental Figure 1:**
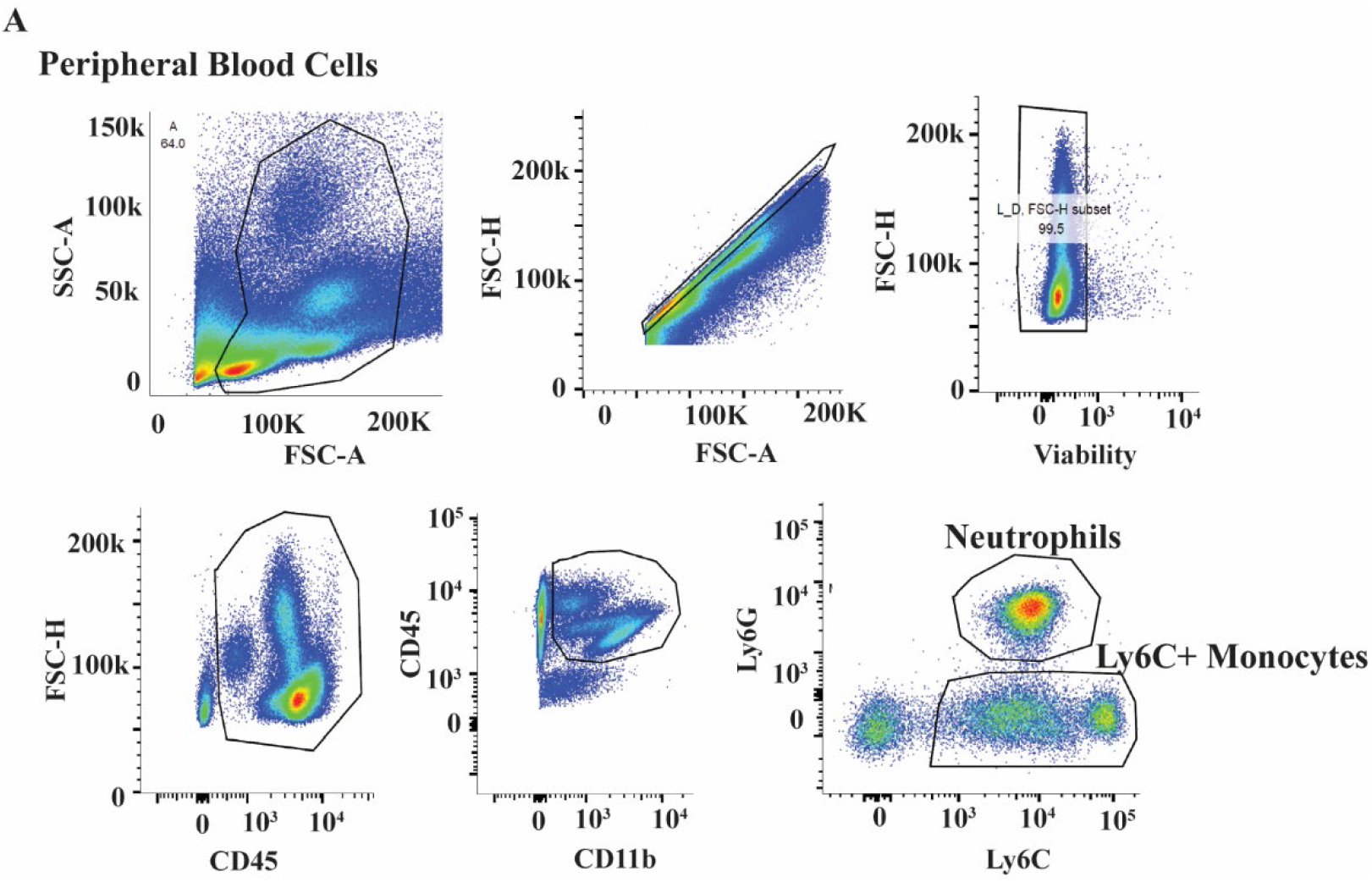
Gating schema for peripheral blood flow cytometry.

**Supplemental Figure 2:**
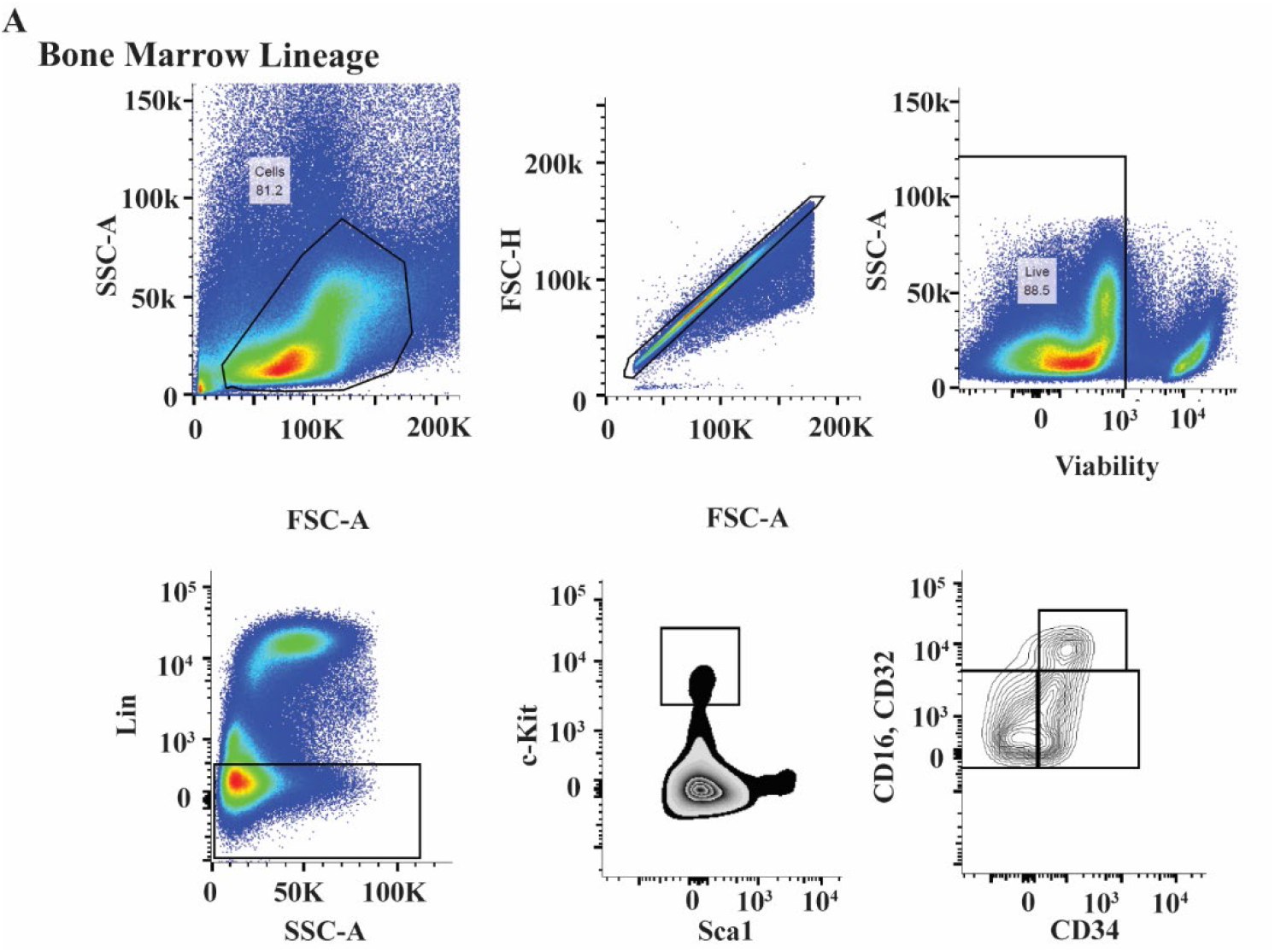
Gating schema for bone marrow blood flow cytometry.

## Notes

### Competing Interest Statement

The authors have declared no competing interest.

## References

1. De Angelis, K., et al., Sympathetic overactivity precedes metabolic dysfunction in a fructose model of glucose intolerance in mice. Am J Physiol Regul Integr Comp Physiol, 2012. 302(8): p. R950–7.

2. Garris, D.R., Developmental and regional changes in brain norepinephrine levels in diabetic C57BL/KsJ mice: effects of estradiol and progesterone. Brain Res Dev Brain Res, 1995. 89(2): p. 314–9.

3. Thaler, J.P., et al., Obesity is associated with hypothalamic injury in rodents and humans. J Clin Invest, 2012. 122(1): p. 153–62.

4. Ding, L., et al., Adipose afferent reflex is enhanced by TNFalpha in paraventricular nucleus through NADPH oxidase-dependent ROS generation in obesity-related hypertensive rats. J Transl Med, 2019. 17(1): p. 256.

5. Mi, Y., et al., Role of microglia M1/M2 polarisation in the paraventricular nucleus: New insight into the development of stress-induced hypertension in rats. Auton Neurosci, 2018. 213: p. 71–80.

6. Hu, P., et al., Loss of survival factors and activation of inflammatory cascades in brain sympathetic centers in type 1 diabetic mice. Am J Physiol Endocrinol Metab, 2015. 308(8): p. E688–98.

7. Sato, I., et al., Peripherally administered baclofen reduced food intake and body weight in db/db as well as diet-induced obese mice. FEBS Lett, 2007. 581(25): p. 4857–64.

8. Fenton, C., G.M. Keating, and K.A. Lyseng-Williamson, Moxonidine: a review of its use in essential hypertension. Drugs, 2006. 66(4): p. 477–96.

9. Thek, K.R., et al., Extensive Inhibitory Gating of Viscerosensory Signals by a Sparse Network of Somatostatin Neurons. J Neurosci, 2019. 39(41): p. 8038–8050.

10. Boehm, S. and S. Huck, A somatostatin receptor inhibits noradrenaline release from chick sympathetic neurons through pertussis toxin-sensitive mechanisms: comparison with the action of alpha 2-adrenoceptors. Neuroscience, 1996. 73(2): p. 595–604.

11. Ikeda, S.R. and G.G. Schofield, Somatostatin blocks a calcium current in rat sympathetic ganglion neurones. J Physiol, 1989. 409: p. 221–40.

12. Mantyh, P.W., et al., Receptor binding sites for cholecystokinin, galanin, somatostatin, substance P and vasoactive intestinal polypeptide in sympathetic ganglia. Neuroscience, 1992. 46(3): p. 739–54.

13. Maynard, K.I., V.L. Saville, and G. Burnstock, Somatostatin modulates vascular sympathetic neurotransmission in the rabbit ear artery. Eur J Pharmacol, 1991. 196(2): p. 125–31.

14. Shapiro, M.S. and B. Hille, Substance P and somatostatin inhibit calcium channels in rat sympathetic neurons via different G protein pathways. Neuron, 1993. 10(1): p. 11–20.

15. Stengel, A., J. Rivier, and Y. Tache, Central actions of somatostatin-28 and oligosomatostatin agonists to prevent components of the endocrine, autonomic and visceral responses to stress through interaction with different somatostatin receptor subtypes. Curr Pharm Des, 2013. 19(1): p. 98–105.

16. Bowman, B.R., et al., Somatostatin 2 Receptors in the Spinal Cord Tonically Restrain Thermogenic, Cardiac and Other Sympathetic Outflows. Front Neurosci, 2019. 13: p. 121.

17. Bhatwadekar, A.D., et al., Hematopoietic stem/progenitor involvement in retinal microvascular repair during diabetes: Implications for bone marrow rejuvenation. Vision Res, 2017. 139: p. 211–220.

18. Karalis, K., et al., Somatostatin analogues suppress the inflammatory reaction in vivo. J Clin Invest, 1994. 93(5): p. 2000–6.

19. Peluso, G., et al., Modulation of cytokine production in activated human monocytes by somatostatin. Neuropeptides, 1996. 30(5): p. 443–51.

20. Markovics, A., et al., Comparison of the anti-inflammatory and anti-nociceptive effects of cortistatin-14 and somatostatin-14 in distinct in vitro and in vivo model systems. J Mol Neurosci, 2012. 46(1): p. 40–50.

21. Mulak, A., et al., Selective agonists of somatostatin receptor subtype 1 or 2 injected peripherally induce antihyperalgesic effect in two models of visceral hypersensitivity in mice. Peptides, 2015. 63: p. 71–80.

22. Basivireddy, J., et al., Somatostatin preserved blood brain barrier against cytokine induced alterations: possible role in multiple sclerosis. Biochem Pharmacol, 2013. 86(4): p. 497–507.

23. Dello Russo, C., et al., Diverging effects of cortistatin and somatostatin on the production and release of prostanoids from rat cortical microglia and astrocytes. J Neuroimmunol, 2009. 213(1-2): p. 78–83.

24. Grinshpun, J., L. Tveria, and S. Fleisher-Berkovich, Differential regulation of prostaglandin synthesis in neonatal rat microglia and astrocytes by somatostatin. Eur J Pharmacol, 2008. 584(2-3): p. 312–7.

25. Dror, N., et al., Inhibitory effect of somatostatin on prostaglandin E2 synthesis by primary neonatal rat glial cells. Regul Pept, 2008. 150(1-3): p. 21–5.

26. Felten, D.L., et al., Noradrenergic and peptidergic innervation of lymphoid tissue. J Immunol, 1985. 135(2 Suppl): p. 755s–765s.

27. Hu, P., et al., CNS inflammation and bone marrow neuropathy in type 1 diabetes. Am J Pathol, 2013. 183(5): p. 1608–20.

28. Vasamsetti, S.B., et al., Sympathetic Neuronal Activation Triggers Myeloid Progenitor Proliferation and Differentiation. Immunity, 2018. 49(1): p. 93–106 e7.

29. Elsaafien, K., et al., Chemoattraction and Recruitment of Activated Immune Cells, Central Autonomic Control, and Blood Pressure Regulation. Front Physiol, 2019. 10: p. 984.

30. Young, J.B., J. Weiss, and N. Boufath, Effects of dietary monosaccharides on sympathetic nervous system activity in adipose tissues of male rats. Diabetes, 2004. 53(5): p. 1271–8.

31. Chaves, V.E., et al., Glyceroneogenesis is reduced and glucose uptake is increased in adipose tissue from cafeteria diet-fed rats independently of tissue sympathetic innervation. J Nutr, 2006. 136(10): p. 2475–80.

32. Dahlman, I., M. Ryden, and P. Arner, Family history of diabetes is associated with enhanced adipose lipolysis: Evidence for the implication of epigenetic factors. Diabetes Metab, 2018. 44(2): p. 155–159.

33. Arner, P., et al., Weight Gain and Impaired Glucose Metabolism in Women Are Predicted by Inefficient Subcutaneous Fat Cell Lipolysis. Cell Metab, 2018. 28(1): p. 45–54 e3.

34. Ryden, M. and P. Arner, Subcutaneous Adipocyte Lipolysis Contributes to Circulating Lipid Levels. Arterioscler Thromb Vasc Biol, 2017. 37(9): p. 1782–1787.

35. Ding, L., et al., Reduced lipolysis response to adipose afferent reflex involved in impaired activation of adrenoceptor-cAMP-PKA-hormone sensitive lipase pathway in obesity. Sci Rep, 2016. 6: p. 34374.

36. Mowers, J., et al., Inflammation produces catecholamine resistance in obesity via activation of PDE3B by the protein kinases IKKepsilon and TBK1. Elife, 2013. 2: p. e01119.

37. Reynisdottir, S., et al., Catecholamine resistance in fat cells of women with upper-body obesity due to decreased expression of beta 2-adrenoceptors. Diabetologia, 1994. 37(4): p. 428–35.

38. Hoogland, I.C., et al., Systemic inflammation and microglial activation: systematic review of animal experiments. J Neuroinflammation, 2015. 12: p. 114.

39. Chung, L., A Brief Introduction to the Transduction of Neural Activity into Fos Signal. Dev Reprod, 2015. 19(2): p. 61–7.

40. Nguyen, N.L., et al., Central sympathetic innervations to visceral and subcutaneous white adipose tissue. Am J Physiol Regul Integr Comp Physiol, 2014. 306(6): p. R375–86.

41. Item, F. and D. Konrad, Visceral fat and metabolic inflammation: the portal theory revisited. Obes Rev, 2012. 13 Suppl 2: p. 30–9.

42. Rytka, J.M., et al., The portal theory supported by venous drainage-selective fat transplantation. Diabetes, 2011. 60(1): p. 56–63.

43. Bartness, T.J., et al., Neural innervation of white adipose tissue and the control of lipolysis. Front Neuroendocrinol, 2014. 35(4): p. 473–93.

44. Souza, S.C., et al., Overexpression of perilipin A and B blocks the ability of tumor necrosis factor alpha to increase lipolysis in 3T3-L1 adipocytes. J Biol Chem, 1998. 273(38): p. 24665–9.

45. Holland, W.L., et al., Inducible overexpression of adiponectin receptors highlight the roles of adiponectin-induced ceramidase signaling in lipid and glucose homeostasis. Mol Metab, 2017. 6(3): p. 267–275.

46. Ohashi, K., et al., Adiponectin promotes macrophage polarization toward an anti-inflammatory phenotype. J Biol Chem, 2010. 285(9): p. 6153–60.

47. Saxton, S.N., et al., Role of Sympathetic Nerves and Adipocyte Catecholamine Uptake in the Vasorelaxant Function of Perivascular Adipose Tissue. Arterioscler Thromb Vasc Biol, 2018. 38(4): p. 880–891.

48. Duan, Y., et al., Loss of Angiotensin-Converting Enzyme 2 Exacerbates Diabetic Retinopathy by Promoting Bone Marrow Dysfunction. Stem Cells, 2018. 36(9): p. 1430–1440.

49. Nagareddy, P.R., et al., Hyperglycemia promotes myelopoiesis and impairs the resolution of atherosclerosis. Cell Metab, 2013. 17(5): p. 695–708.

50. Beli, E., et al., CX3CR1 deficiency accelerates the development of retinopathy in a rodent model of type 1 diabetes. J Mol Med (Berl), 2016. 94(11): p. 1255–1265.

51. Barman, P.K., N. Urao, and T.J. Koh, Diabetes induces myeloid bias in bone marrow progenitors associated with enhanced wound macrophage accumulation and impaired healing. J Pathol, 2019. 249(4): p. 435–446.

52. Bhatwadekar, A.D., et al., Ataxia Telangiectasia Mutated Dysregulation Results in Diabetic Retinopathy. Stem Cells, 2016. 34(2): p. 405–17.

53. Luu, C.D., et al., Correlation between retinal oscillatory potentials and retinal vascular caliber in type 2 diabetes. Invest Ophthalmol Vis Sci, 2010. 51(1): p. 482–6.

54. Rajagopal, R., et al., Functional Deficits Precede Structural Lesions in Mice With High-Fat Diet-Induced Diabetic Retinopathy. Diabetes, 2016. 65(4): p. 1072–84.

55. Musovic, S. and C.S. Olofsson, Adrenergic stimulation of adiponectin secretion in visceral mouse adipocytes is blunted in high-fat diet induced obesity. Sci Rep, 2019. 9(1): p. 10680.

56. Qi, Z. and S. Ding, Obesity-associated sympathetic overactivity in children and adolescents: the role of catecholamine resistance in lipid metabolism. J Pediatr Endocrinol Metab, 2016. 29(2): p. 113–25.

57. Busik, J.V., et al., Diabetic retinopathy is associated with bone marrow neuropathy and a depressed peripheral clock. J Exp Med, 2009. 206(13): p. 2897–906.

58. Clegg, D.J., et al., Consumption of a high-fat diet induces central insulin resistance independent of adiposity. Physiol Behav, 2011. 103(1): p. 10–6.

59. Moraes, J.C., et al., High-fat diet induces apoptosis of hypothalamic neurons. PLoS One, 2009. 4(4): p. e5045.

60. McNay, D.E., et al., Remodeling of the arcuate nucleus energy-balance circuit is inhibited in obese mice. J Clin Invest, 2012. 122(1): p. 142–52.

61. Weissmann, L., et al., IKKepsilon is key to induction of insulin resistance in the hypothalamus, and its inhibition reverses obesity. Diabetes, 2014. 63(10): p. 3334–45.

62. Garris, D.R., R.L. West, and D.L. Coleman, Morphometric analysis of medial basal hypothalamic neuronal degeneration in diabetes (db/db) mutant C57BL/KsJ mice: relation to age and hyperglycemia. Brain Res, 1985. 352(2): p. 161–8.

63. Martyniuk, C.J., et al., Genetic ablation of bone marrow beta-adrenergic receptors in mice modulates miRNA-transcriptome networks of neuroinflammation in the paraventricular nucleus. Physiol Genomics, 2020. 52(4): p. 169–177.

64. Mickelsen, L.E., et al., Single-cell transcriptomic analysis of the lateral hypothalamic area reveals molecularly distinct populations of inhibitory and excitatory neurons. Nat Neurosci, 2019. 22(4): p. 642–656.

65. Chamessian, A., et al., Transcriptional Profiling of Somatostatin Interneurons in the Spinal Dorsal Horn. Sci Rep, 2018. 8(1): p. 6809.

66. Han, C., M.W. Rice, and D. Cai, Neuroinflammatory and autonomic mechanisms in diabetes and hypertension. Am J Physiol Endocrinol Metab, 2016. 311(1): p. E32–41.

67. Li, M., et al., TNF-alpha Upregulates IKKepsilon Expression via the Lin28B/let-7a Pathway to Induce Catecholamine Resistance in Adipocytes. Obesity (Silver Spring), 2019. 27(5): p. 767–776.

68. Santos, M.P., et al., A low-protein, high-carbohydrate diet increases fatty acid uptake and reduces norepinephrine-induced lipolysis in rat retroperitoneal white adipose tissue. Lipids, 2012. 47(3): p. 279–89.

69. Sipe, L.M., et al., Differential sympathetic outflow to adipose depots is required for visceral fat loss in response to calorie restriction. Nutr Diabetes, 2017. 7(4): p. e260.

70. Vieira, C.P., et al., Selective LXR agonist DMHCA corrects retinal and bone marrow dysfunction in type 2 diabetes. JCI Insight, 2020. 5(13).

71. Weeke, J., et al., A randomized comparison of intranasal and injectable octreotide administration in patients with acromegaly. J Clin Endocrinol Metab, 1992. 75(1): p. 163–9.

72. Anaclet, C., et al., Genetic Activation, Inactivation, and Deletion Reveal a Limited And Nuanced Role for Somatostatin-Containing Basal Forebrain Neurons in Behavioral State Control. J Neurosci, 2018. 38(22): p. 5168–5181.

73. Todd, W.D., et al., Suprachiasmatic VIP neurons are required for normal circadian rhythmicity and comprised of molecularly distinct subpopulations. Nat Commun, 2020. 11(1): p. 4410.

